# Hypoxia Depletes Contaminating CD45^+^ Hematopoietic Cells from Bone Marrow Stromal Cell (BMSC) Cultures: Methods for BMSC Culture Purification

**DOI:** 10.1101/2020.04.29.068031

**Authors:** Wendi Guo, Kassandra V. Spiller, Jackie Tang, Courtney M. Karner, Matthew J. Hilton, Colleen Wu

## Abstract

Culture expanded bone marrow stromal cells (BMSCs) are easily isolated, can be grown rapidly *en masse,* and contain both skeletal stem cells (SSCs) and multipotent mesenchymal progenitors (MMPs). Despite this functional heterogeneity, BMSC cultures continue to be utilized for many applications due to the lack of definitive and universally accepted markers to prospectively identify and purify SSCs. Isolation of BMSCs is widely based on their adherence to tissue culture plastic; however, this method poses unique challenges when working with murine models. In particular, hematopoietic cell contamination is a significant impediment during the isolation or murine BMSCs. Several strategies to reduce hematopoietic contamination have been developed with varying levels of success. In this study, we found that up to 60% of cells isolated in standard BMSC cultures expressed the hematopoietic marker CD45. Remarkably, when cultured at a physiological oxygen tension of 1% O_2_, there was a 10-fold reduction in hematopoietic cells associated with a concomitant increase in PDGFRα^+^ expressing stromal cells. This finding was due, in part, to a differential response of the two populations to low oxygen tension, or hypoxia. Specifically, in standard tissue culture conditions of 21% O_2_, CD45^+^ hematopoietic cells showed increased proliferation coupled with no observable changes in cell death when compared to their counterparts grown at 1% O_2_. In contrast, PDGFRα^+^ stromal cells responded to hypoxia by increasing proliferation and exhibiting a 10-fold decrease in cell death. In summary, we describe a simple and reliable method exploiting the divergent biological response of hematopoietic and stromal cells to hypoxia to significantly increase the PDGFRα^+^ stromal cell population in murine BMSC cultures.

## INTRODUCTION

The skeleton houses SSCs which arise in post-natal bone and are defined by their *in vivo* ability to self-renew and to differentiate into hematopoietic supporting stroma, osteogenic, chondrogenic and adipogenic lineages[1–3]. Currently, there remains a lack of definitive cell surface markers to identify and/or prospectively isolate a homogenous population of SSCs [4–6]. In lieu of this, BMSCs have been shown to support hematopoiesis and contribute to all functions of the skeleton. BMSCs were first isolated based on their rapid adherence to tissue culture (TC) plastic [7]. At non-clonal density, they can be quickly expanded and are capable of differentiating into distinct mesodermal lineages *in vitro* demonstrating the presence of MMPs [8]. When plated at clonogenic density, some BMSCs exhibit density independent growth, forming discrete colonies derived from single cells which are referred to as colony forming units-fibroblastic (CFU-F) [9–11]. Upon transplantation, some CFU-Fs have the capacity for *in vivo* multipotent differentiation and self-renewal, the defining features of SSCs [12, 13]. For these reasons, BMSCs serve as a powerful tool for cell-based therapies to treat musculoskeletal disorders by repairing damaged mesenchymal tissues. Moreover, with the breadth of transgenic and genetically engineered mouse models, BMSCs also serve as an invaluable means to harness information regarding the cellular and molecular mechanisms regulating MMP and SSC function. Importantly, the successful use of BMSCs for cell-based therapies and meaningful interpretation of genetic studies is dependent upon the ability to isolate pure populations of mesenchymal cells.

In humans, preferential attachment to TC plastic can be used with success for the isolation of mesenchymal populations; however, for murine models, this technique presents several challenges [14, 15]. Most notably, this includes high hematopoietic contamination because murine hematopoietic cells can directly adhere to TC plastic and both hematopoietic stem cells (HSCs) and their differentiated progeny can attach directly to stromal cells [16, 17]. Moreover, stromal cells have the capacity to support *in vitro* granulopoiesis in the absence of exogenous cytokines [18]. Collectively, these factors contribute to the persistent contamination of hematopoietic cells in murine BMSC cultures even after prolonged culturing [16, 19].

Several methodologies such as subculturing, immunodepletion, and low-density seeding have been tested to eliminate hematopoietic cells from BMSC cultures; however, these techniques have only achieved limited success. While frequent and prolonged subculturing improves purity, it also results in a replicative senescent phenotype with the possibility of altered differentiation potential [20, 21]. Furthermore, this method increases the potential of selecting for clones carrying inactivating mutations in p53, thereby increasing the likelihood of cellular transformation [22, 23]. Immunodepletion can result in a limited proliferative capacity of BMSCs [24]. Because only a small number of cells are recovered, initiation of cultures at low densities is prohibitively time consuming where rapid expansion of multipotent mesenchymal stromal cells is required [25]. Taken together, these limitations demonstrate that hematopoietic contamination continues to serve as a significant impediment to the isolation of BMSCs based on plastic adherence [26].

The bone microenvironment is characterized by vascular heterogeneity associated with regional areas of hypoxia [27–29]. Under hypoxic conditions, cells upregulate the hypoxia inducible factor (HIF) signaling pathway to drive expression of genes which facilitate oxygen delivery and cellular adaptation to low oxygen [30]. Importantly, activation of HIF target genes is highly context dependent with cellular responses exquisitely sensitive to oxygen concentration [31]. Imaging analysis of HSC and SSC cell surface markers demonstrate that these cells are localized to perivascular regions where direct and indirect measurements reveal oxygen tensions ranging from 4 – 1% [1, 32]. While both human and murine BMSCs demonstrate improved proliferation capacity at 5% O_2_, the differential response of mesenchymal stromal cells and hematopoietic progenitors to physiologic oxygen tensions in the bone marrow is less well defined [22]. For these reasons, we aimed to determine the impact of culturing BMSCs under physiologic hypoxic conditions of 1% O_2_. Importantly, we discovered that 1% O_2_ strikingly reduced the percentage of CD45 expressing hematopoietic cells and this reduction remained consistent over time. Here, we report an easy and reliable method by which hematopoietic cells can be selectively eliminated while simultaneously enriching for stromal cells during BMSC preparations.

## RESULTS

### The Majority of Bone Marrow Cells Isolated Based on Adherence to Tissue Culture Plastic Express the Hematopoietic Marker CD45

We isolated BMSCs based on the rapid adherence of cells to TC plastic when collected from bone marrow flushes. We noted that high density BMSC (1.3 x 10^5^ cm^2^) cultures were phenotypically heterogeneous when incubated under 21% O_2_, with many cells exhibiting small rounded morphology indicative of monocytes (**Figure 1A, C**). Given this morphological heterogeneity, we sought to determine the proportion of presumptive MMP and SSCs in our cultures. For this purpose, we first evaluated the expression of a known bone marrow stromal cell marker which enriches for SSCs, the leptin receptor (LepR). LepR marks a population of bone marrow stromal cells which arise perinatally and contribute to osteoblast, chondrocyte and adipocyte lineages *in vivo* [33]. Flow cytometry and imaging analysis of BMSCs isolated from *LeprCre;Rosa26^tdTomato/+^* conditional reporter mice revealed only a small proportion of cells expressed tdTomato (Td/Tm) (**Figure 1A**). To confirm our analysis we evaluated the expression of PDGFRα, an early mesenchymal marker which is broadly expressed in the bone marrow stroma, highly expressed in CFU-Fs, and multipotent when subjected to *in vitro* differentiation assays [34]. Consistent with other reports, LepR^+^ bone marrow stromal cells uniformly express PDGFRα, supporting the concept that bone marrow stromal cells express both PDGFRα and LepR (**Figure 1A**). Notably, the proportion of tdTomato^+^, PDGFRα^+^ or tdTomato^+^;PDGFRα^+^, double positive cells constituted only a minority of the total cell population when using standard BMSC isolation methods (**Figure 1A)**.

**Figure 1.**
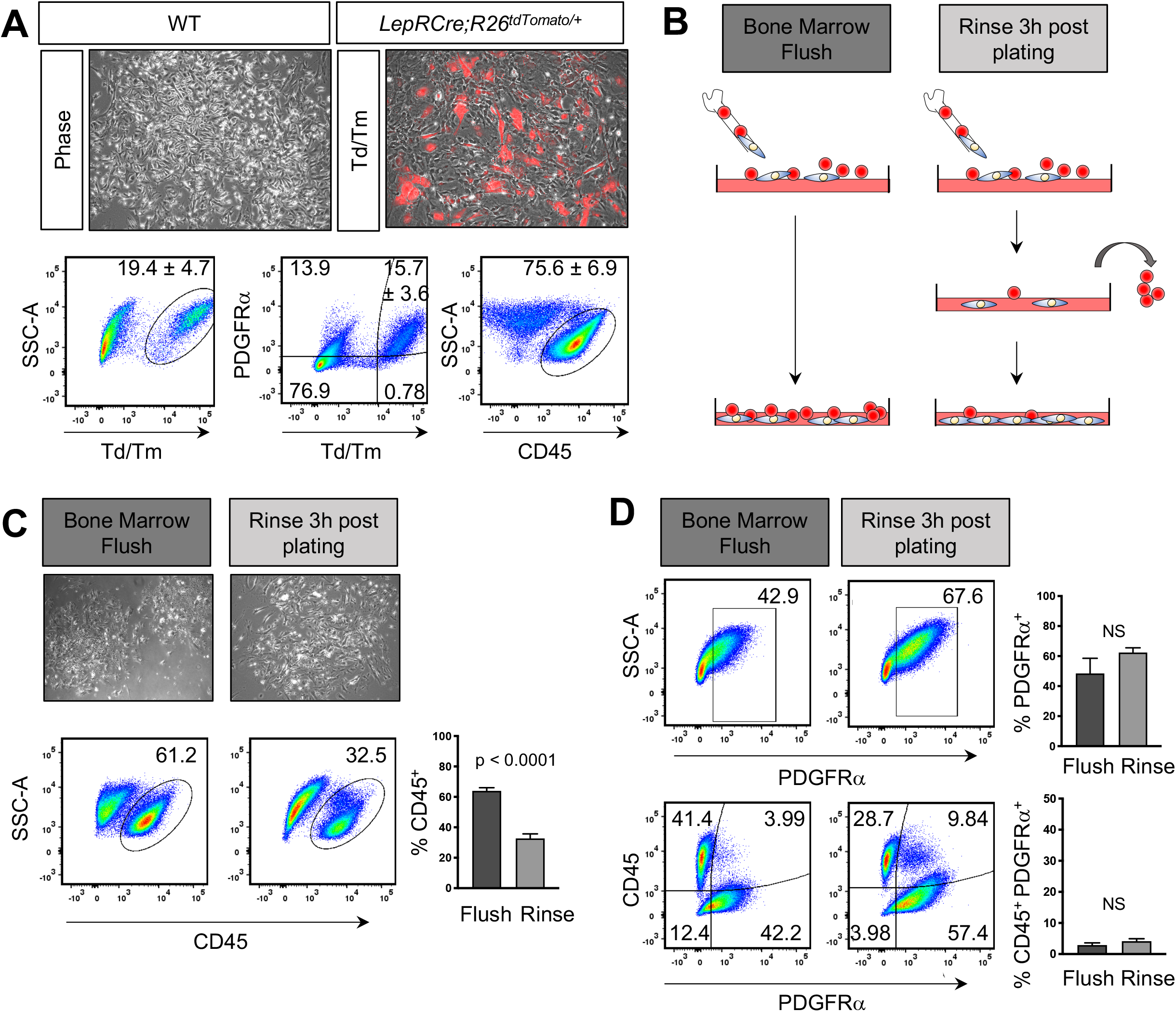
CD45^+^ cells constitute the majority of cells in BMSC cultures. **A)** Phase contrast and overlay of immunofluorescent and phase contrast images of BMSCs isolated from *LepRCre;Rosa26^tdTomato/+^* mice. Red depicts cells in which CRE recombinase was active. Representative flow analysis plots and quantification of the percentage of CD45^+^, tdTomato^+^ (Td/Tm), and PDGFRα^+^;tdTomato^+^ double positive cells isolated from *LepRCre;Rosa26^tdTomato/+^* mice. **B)** Schematic representation of BMSC isolation methods. **C)** Phase contrast images of passage 0 BMSCs after 7 days in culture. Representative flow analysis plots and quantification of the percentage **C)** CD45^+^, **D)** PDGFRα^+^, and PDGFRα^+^;CD45^+^ double positive cells in WT BMSCs. Gates for positive cells denoted by outlined areas. Each graph represents an individual trial using BMSCs isolated from 2-3 month male and female mice. n≥3, unpaired two tailed t-tests, significance p < 0.05.

Given the low percentage of MMP and SSCs, we next sought to determine the remaining cellular constituents within our cultures. Remarkably, using a gating strategy in which isotype controls acted as a negative reference, flow cytometry analysis revealed that the majority of cells in BMSCS cultures express the hematopoietic cell surface marker CD45 63.98 ± 2.03% (**Figure S1A**, **1A, B**). Given the high hematopoietic contamination present in these cultures, we sought to determine if vigorously rinsing dishes 3 hours after plating would reduce the number of CD45^+^ cells (**Figure 1B**) [10]. While the majority removed cells were CD45^+^ (**Figure S1B);** despite vigorous rinsing, a high percentage of contaminating CD45^+^ cells remained in our BMSC cultures (**Figure 1C**). Strikingly, in unrinsed samples, only 48.43 ± 10.0% of cells expressed PDGFRα (**Figure 1D)**. Vigorous rinsing after plating led to an increasing trend of PDGFRα^+^ cells; however, these findings were not statistically significant suggesting that rinsing may remove only CD45 expressing cells that take longer to adhere to TC plastic or to stromal cells and not enrich for PDGFRα^+^ cells (**Figure 1D, S1B**). Notably, PDGFRα expression was primarily restricted to CD45^-^ cells (**Figure 1D)**. Taken together, these data suggest that isolation of BMSCs with conventional techniques leads to a high contamination of CD45^+^ hematopoietic cells coupled with moderately low numbers of mesenchymal stromal cells. These findings highlight the need for methods to increase the purity of stromal cells during BMSC isolation.

### Hypoxia Reduces CD45^+^ Cells While Increasing PDGFRα^1^’ Cells Present in BMSC Cultures

Given the physiological relevance of hypoxia in regulating cell function within the bone microenvironment, we sought to determine if this environmental factor could influence the cellular composition of BMSC preparations. To test this, BMSCs were isolated, rinsed and grown either at atmospheric (21%; normoxia) or physiologic (1%; hypoxia) oxygen tensions. In contrast to normoxic cultures which remained heterogeneous, phase microscopy revealed that hypoxic cultures consisted of a homogeneous population of large bipolar spindle shaped cells (**Figure 2A**). Interestingly, flow analysis demonstrated that larger cells constituted the majority of CD45^-^, PDGFRα^+^ stromal cells (**Figure S2**). Consistent with the observed changes in cellular morphology, cultures grown in hypoxic conditions showed a significant reduction in CD45^+^ cells (45.12 ± 4.44 % vs 5.78 ± 0.95 %) coupled with a substantial increase in PDGFRα^+^ cells (48.90% ± 5.73 % vs 81.48 ± 2.86 %) when compared to cells grown in normoxic conditions (**Figure 2B**). Supporting these findings, BMSCs isolated from *LepRCre;R¤sa26^tdTomato/+^* mice and grown at 1% O_2_ also showed a significant decrease in CD45^+^ cells which was associated with an increase in both tdTomato^+^, PDGFRα^+^ and tdTomato^+^;PDGFRα^+^ double positive cells (**Figure 2C, D**). Taken together, these findings demonstrate that modifying culture conditions by decreasing oxygen levels to 1% profoundly diminishes the percentage of hematopoietic contaminants while enriching for PDGFRα^+^/LepRCre^+^ derived stromal cells in BMSC cultures.

**Figure 2.**
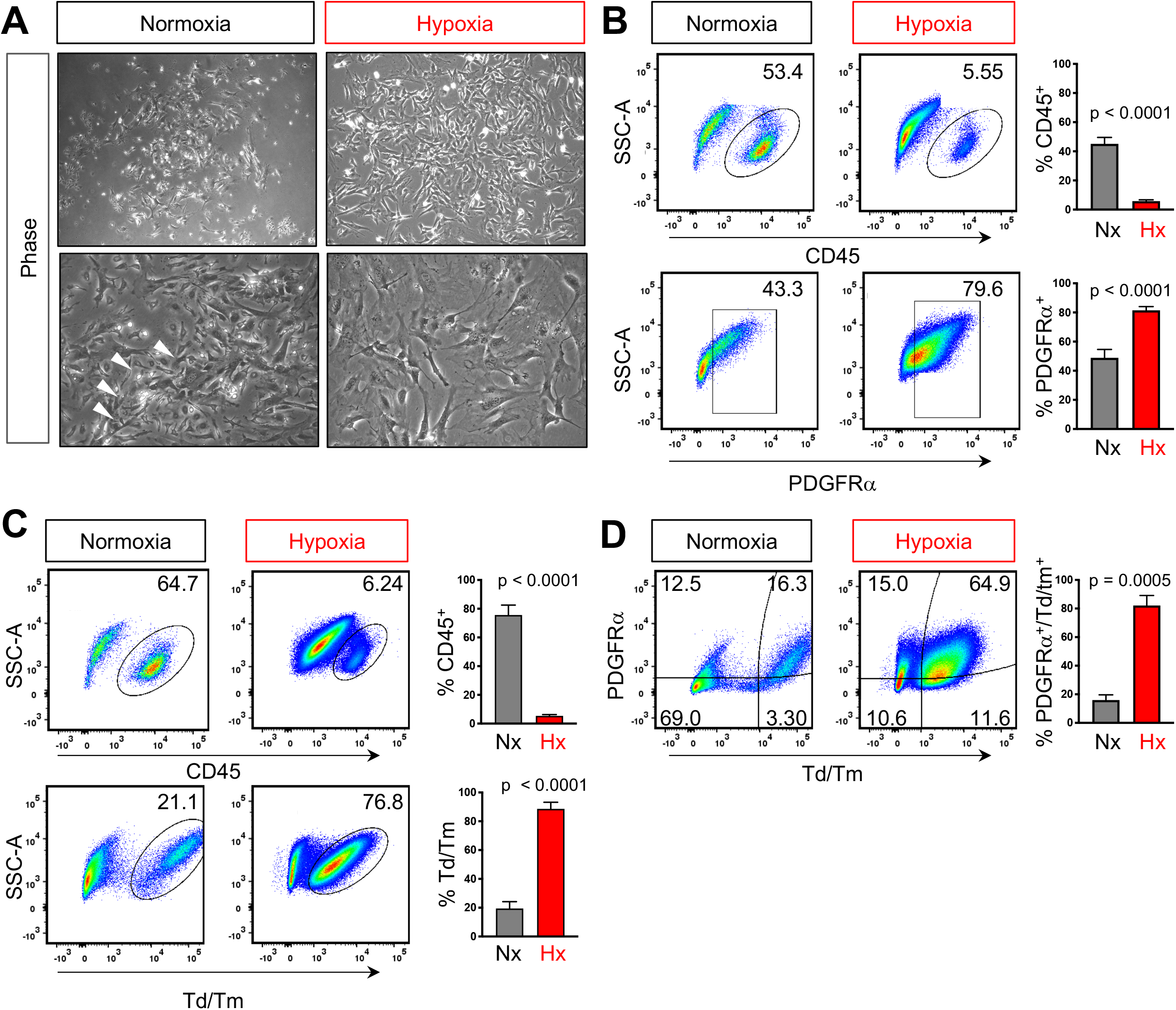
Hypoxia depletes CD45^+^ cells in BMSC cultures. **A)** Phase contrast images of passage 0 BMSCs after 7 days in either 21% O_2_ (normoxia, Nx) or 1% O_2_ (hypoxia, Hx). White arrowheads denote small rounded cells. Representative flow analysis plots and quantification of the percentage of CD45^+^, PDGFRα^+^ and PDGFRα^+^;tdTomato^+^ double positive cells in **B)** WT or **C)** *LepRCre;Rosa26^tdTomato/+^* BMSCs cultured in either normoxia or hypoxia. Gates for positive cells denoted by outlined areas. Each graph represents an individual trial using BMSCs isolated from 2-3 month male mice were used for analysis, n≥3, unpaired two tailed t-tests, significance p < 0.05

### Hypoxic Reduction of CD45^+^ Cells Remains Constant Over Time, Passage Number, and Cell Density

To further explore the relationship between hypoxia and the reduction in hematopoietic cells, we first sought to determine if CD45 expression remained consistent over time in low oxygen. We noted that over a period of seven days, CD45^+^ cells remained consistently high in normoxic cultures but remained below 6% in hypoxic cultures at all time points (**Figure 3A**). Moreover, after two rounds of trypsinization, CD45^+^ cells continued to persist in normoxic cultures but were reduced to 1.36 ± 0.19% of cells in hypoxic cultures (**Figure 3B**). While there was an increase in PDGFRα cells cultured under normoxic conditions in passage 2 (P2) when compared to passage 1 (P1), this was likely due to an increase in CD45^+^;PDGFRα^+^ double positive cells, suggesting that over time in normoxia, CD45^+^ cells increase their expression of the PDGFRα receptor (**Figure 3B**).

**Figure 3.**
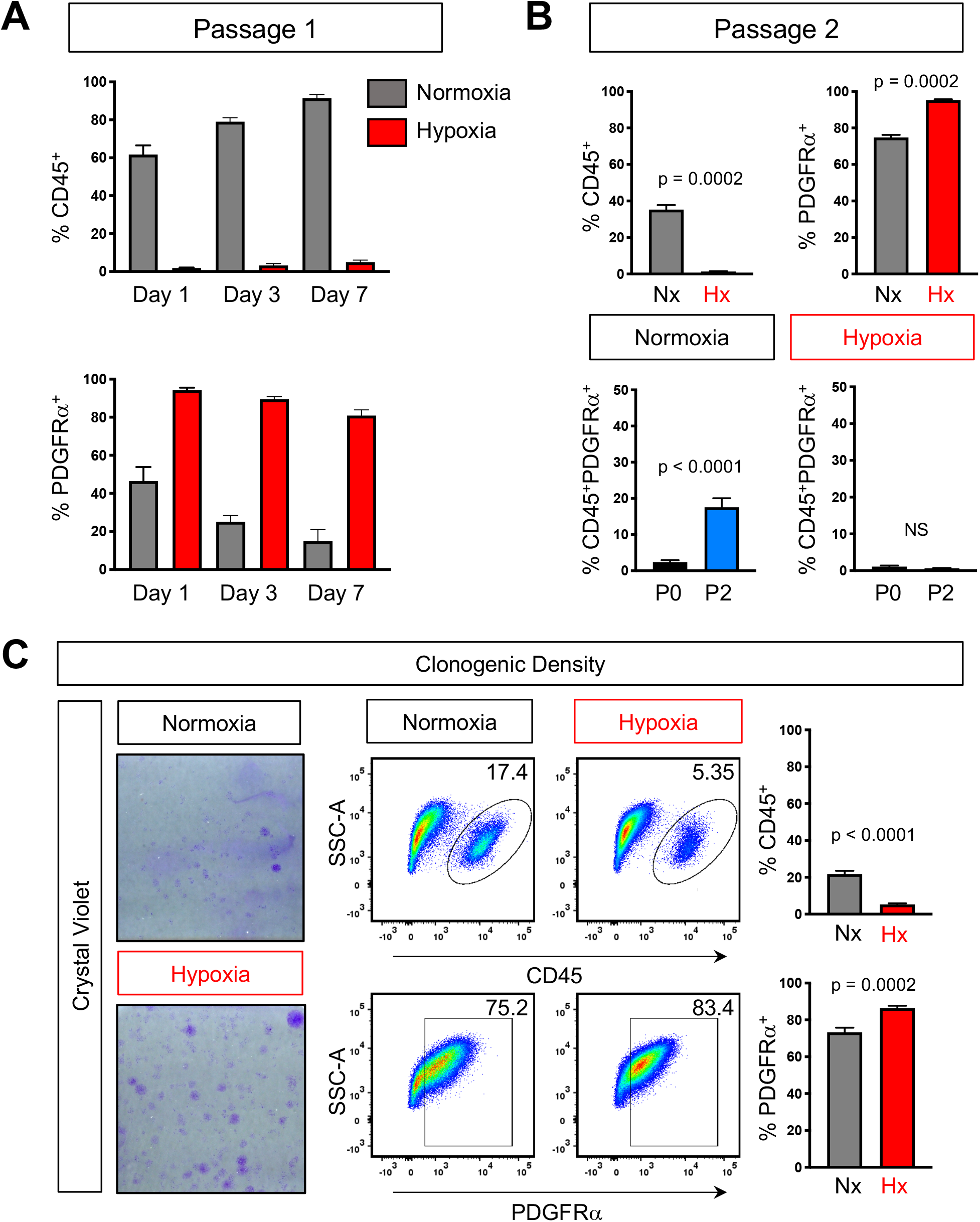
Hypoxic depletion of CD45^+^ cells is maintained over time and passage number. **A)** Quantification of flow analysis showing the percentage of CD45^+^ and PDGFRα cells over 7 days in either normoxia (grey) or hypoxia (red). **B**) Quantification of the percentage of CD45^+^, PDGFRα^+^, and CD45^+^;PDGFRα^+^ double positive cells after two passages in either normoxia or hypoxia. **C)** Crystal violet staining of BMSCs plated at clonogenic density (left) and representative flow analysis plots and quantification of the percentage of CD45^+^ and PDGFRα^+^ (right). Each graph represents an individual trial using BMSCs isolated from 2-3 month male mice were used for analysis, n≥3, unpaired two tailed t-tests, significance p < 0.05

Culturing cells at low density has been shown to reduce hematopoietic contaminates [25]. As such, we next sought to determine if hypoxia would also enrich the cellular composition of stromal cells in BMSC cultures grown at clonogenic density. Consistent with previous reports, when cells were plated at 6 x 10^5^ cells/cm^2^ with rinsing, crystal violet staining highlighted the presence of discrete colonies, derived from single cells (**Figure 3C**). Importantly, hypoxia increased both the total number of colonies and their overall size (**Figure 3C**). Accordingly, flow cytometry analysis revealed a significant reduction in the proportion of hematopoietic cells when compared to high density BMSC cultures, however, 21.82% ± 1.73 % of cells were still of hematopoietic origin. In contrast, when grown in hypoxia, 5.04 % ± 0.53 % of cells expressed CD45 (**Figure 3C**). Additionally, the percentage of PDGFRα^+^ cells in hypoxia was also increased relative to normoxia (**Figure 3C)**. While at clonogenic density, the number of CD45^+^ cells are fewer than when plated at high density cultures; importantly, hypoxia is still able to reduce hematopoietic cells in these conditions. Taken together, our data demonstrates that BMSCs grown under physiological hypoxia results in a dramatic decrease of contaminating CD45^+^ hematopoietic cells while increasing PDGFRα^+^ cells which is consistent over time, passage number and cell density.

### Hypoxia Reduces Proliferation of CD45^+^ Cells While Inhibiting Apoptosis of PDGFRα^+^ Cells

The differential effect of hypoxia on hematopoietic and stromal components isolated during BMSC preparations has not been previously reported. To determine the cellular mechanisms regulating hypoxia dependent reduction of CD45^+^ cells in our cultures, we first assessed proliferation by incubating cells with EdU. Flow cytometry analysis revealed CD45^+^ cells exhibit increased EdU incorporation in normoxia when compared to hypoxia (**Figure 4A,C**). Conversely, in normoxic conditions we noted a decrease in PDGFRα^+^;EdU^+^ double positive cells when compared to hypoxic conditions (**Figure 4B,C**). Importantly, in normoxia, CD45^+^ cells had higher levels of EdU incorporation when compared to PDGFRα^+^ cells; while in hypoxia, CD45^+^ cells were less proliferative than PDGFRα^+^ cells (**Figure 4C**). We next evaluated apoptosis by performing TUNEL staining assays to assess DNA fragmentation. We did not detect statistically significant differences between CD45^+^ cells subjected to varying oxygen tensions (**Figure 4D, F**). However, there was a 10-fold decrease of PDGFRα^+^ cells undergoing programmed cell death in hypoxia when compared to normoxia (**Figure 4E, F)**. Moreover, in normoxia, we detected a 6-fold increase in PDGFRα^+^ cells undergoing apoptosis when compared to CD45^+^ cells (**Figure 4F**) but no differences between the two cell populations were detected under hypoxia. Taken together, these data indicate that CD45^+^ hematopoietic cells are more proliferative while PDGFRα^+^ cells undergo much greater cell death at 21% oxygen. However, in hypoxia, PDGFRα^+^ cells are more proliferative while both populations undergo cell death at the same low rate. Hence, this differential response in proliferation and apoptosis likely contributes to diminished CD45^+^ cells and the associated expansion of PDGFRα^+^ cells at low oxygen tensions.

**Figure 4.**
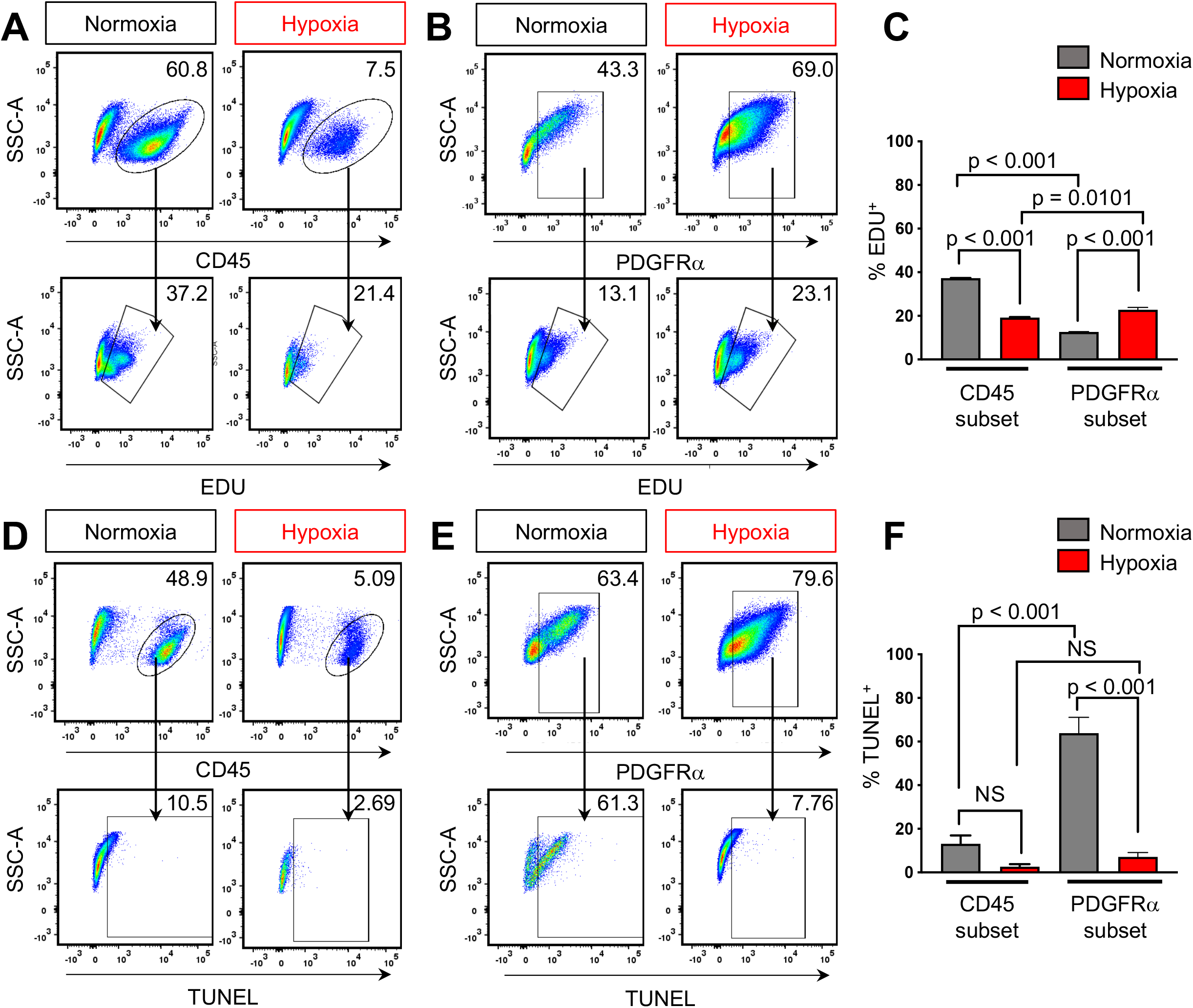
Differential effect of hypoxia on proliferation and apoptosis within CD45 and PDGFRα subsets. **A)** Representative flow analysis plots of CD45^+^ cells or **B)** PDGFRα^+^ and percentage of cells which have incorporated EdU under normoxic or hypoxic conditions **C)** Quantification of flow analysis for CD45^+^;EdU^+^ and PDGFRα^+^;EdU^+^ cells in either normoxic or hypoxic conditions. **D)** Representative flow analysis plots of CD45^+^ cells or **E)** PDGFRα^+^ and percentage of TUNEL^+^ cells under normoxic or hypoxic conditions **F)** Quantification of flow analysis for CD45^+^;TUNEL^+^ and PDGFRα^+^;TUNEL^+^ cells in either normoxic or hypoxic conditions. BMSCs isolated from 2-3 month old male and female mice, n≥3 for each condition. One-way ANOVA, post-hoc Tukey’s multiple comparisons test, significance p < 0.05.

## DISCUSSION

Although physiologic oxygen tensions in the bone microenvironment range from 4% – 1%, tissue culture incubators are standardly set to mimic atmospheric oxygen conditions of 21% [28]. Consistent with other reports, we found at this oxygen tension, the majority of cells in murine BMSC cultures expressed the hematopoietic marker, CD45 [22, 24]. Standard protocols that incorporate a vigorous rinsing step three hours after plating reduced CD45^+^ hematopoietic contaminating cells by ~50%. Here, we demonstrate incubation at a physiological oxygen tension of 1%reduced hematopoietic contaminants by over 90% during murine bone marrow preparations (Figure 2B). Importantly, culturing at 1% oxygen also had the added benefit of selecting for PDGFRα^+^ stromal cells. It is important to note that a previous study culturing BMSCs at 5% O_2_ demonstrated improved murine BMSCs lifespan and differentiation potential; however, CD45 expressing cells remained prevalent, constituting 49% of the total cell population. To reduce hematopoietic contamination, the authors performed serial immunodepletion utilizing antibodies against lineage markers [22, 35]. In contrast, our data indicates culturing BMSCs directly in 1% O_2_ is sufficient to dramatically reduce contaminating CD45^+^ cells to less than 6% of the total cellular population, thus potentially eliminating the need for immunodepletion. Moreover, this method reduces the need for additional passaging of cells required for immunodepletion; thereby decreasing the possibility of cellular transformation or immortalization which is associated with increased passage number [36, 37].

The bone microenvironment provides the supportive niche for HSCs and SSCs while also housing their differentiated progeny. During HSC differentiation and maturation, hematopoietic cells switch from glycolysis to oxidative phosphorylation to allow for cells to respond to increasing energy demands associated with differentiation [38–40]. Hence, despite its high vascular density, the bone marrow microenvironment harbors regions which are strongly hypoxic due in part to the high oxygen consumption by resident hematopoietic cells [27–29]. Indeed, Kroghian models of the bone microenvironment estimate that a three-cell layer of granulocytic progenitors is sufficient to deplete the oxygen of a nearby sinusoid [41]. Accordingly, this differentiation process is associated with an increase in mitochondrial reactive oxygen species (ROS) created during oxidative phosphorylation [42, 43]. We noted in normoxic conditions, CD45 expressing cells exhibited an increase in EdU incorporation when compared to cells grown in hypoxic conditions with no observed differences in TUNEL^+^ staining. It is possible that differentiated hematopoietic cells present in BMSC cultures are inherently resistant to ROS generated at 21% O_2_ due to the metabolic reprogramming that occurs during differentiation. Murine HSCs, are localized to areas of low oxygen tension, are defined by strong retention of the hypoxia marker, pimonidazole, and require HIF-1a for survival and quiescence [44, 45]. As such, we cannot rule out the possibility that these biological characteristics leads to the retainment of HSC in hypoxia thereby contributing to low level of CD45^+^ cells persisting at 1% O_2_. Collectively, the differential response seen between CD45^+^ and PDGFRα^+^ cells suggests that inherent differences in biological response to oxygen can be exploited to enhance purification of BMSC cultures.

Supraphysiologic levels of oxygen can result in a several fold increase in ROS causing extensive damage to DNA leading to genomic instability, cell cycle arrest and cell death [46]. Indeed, high oxygen tension has been shown to inhibit proliferation and survival of both human and murine BMSCs, attributed in part, to p53 dependent mitochondrial ROS production [22]. Consistent with these studies, our data indicates PDGFRα^+^ BMSC are exquisitely sensitive to high oxygen tension, undergoing rapid cell death as indicated by the high percentage of TUNEL positive cells when compared to CD45^+^ hematopoietic cells. BMSC cultured in 1% oxygen were characterized by a 10-fold reduction in programmed cell death when compared to their normoxic counterparts. These data are consistent with studies demonstrating reduced BMSC apoptosis when cultured at 3-5% oxygen [22, 47].

Previous reports in murine models demonstrated that hematopoietic cells are required to provide paracrine factors necessary for stromal cell growth. Specifically, after 8 days of culture, immunodepleted BMSCs isolated from FVB/N and Balb/C mice exhibited limited growth potential relative to non-immunodepleted cells [11, 13, 24]. However, this growth inhibition was not observed in immunodepleted cells in hypoxic conditions [22]. In our study, hypoxic cultures passaged after 7 days showed incorporation of EdU suggesting the retainment of proliferative capacity. Moreover, our studies were performed using BMSCs isolated from C57BL/6 mice which have growth rates much lower than both FVB and Balb/C mice suggesting that this technique could be used to purify and overcome the inherent growth limitations in BMSCs isolated form C57BL/6 stain of mice [48]. However, we cannot rule out the possibility that the low level of CD45^+^ cells present in hypoxic cultures was sufficient to provide paracrine factors for stromal cells.

In summary, several methodologies have been developed for the isolation of more homogeneous populations of BMSCs including prospective isolation using cell surface markers [32], plating at low densities [13], and immunodepletion; however, these protocols have not been widely adopted. Difficulties in purifying BMSCs arise from limited lifespan, the capacity to undergo transformation and high hematopoietic content. In this report, we identified that isolating and culturing BMSCs at 1% O_2_ affords an easy and reproducible method to improve the purity of BMSC cultures that can be universally adopted.

## EXPERIMENTAL PROCEDURES

### Culture of bone marrow-derived stromal cells

Bone marrow stromal cells were isolated from 8-16-week-old male and female C57BL/6J mice (Jackson Laboratory, 000664). All experimental procedures were approved by the Institutional Animal Care and Use Committee at Duke University. Mice were euthanized by CO_2_ asphyxiation and long bones were subsequently extracted. BMSCs were isolated by flushing the bone marrow of the extracted femurs and tibias. Prior to plating, red blood cells were removed using an ammonium chloride-based red blood cell lysis buffer (Sigma-Aldrich, R7757). Cells were cultured in αMEM containing nucleosides (Gibco, 12571-063) supplemented with 20% fetal bovine serum (FBS) (GE Healthcare Life Sciences, SH30071.03), and 1% Penicillin-Streptomycin (P/S) (Gibco, 15140-122) at 37°C in 5% CO_2_ atmosphere. Three hours post-plating, the BMSCs cultures were subjected to a 3x PBS rinse to remove non-adherent cells.

For unrinsed cells, cells were cultured as described above, however, the 3 hours post-plating rinse was not performed. Media was changed 2 days post-plating and cells were grown for 7 days in their respective conditions.

To assess clonogenic densities, cells were cultured and grown for 7 days as described above and seeded at 6 x 10^4^ per cm^2^.

For the passage 0 time point, cells were seeded at a density of 40 million cells per 150 mm dish for normoxic (21% O_2_) cultures and 13 million cells per 100 mm dish for hypoxic cultures prior to rinsing. Hypoxic cell cultures were grown in an InvivO_2_ 400 chamber (Baker Ruskinn) at 1% oxygen tension. Following a 3-hour post-plating rinse, cells were grown for 7 days in their respective conditions.

For passage 1 time points, cells were initially seeded at the densities described in the passage 0 time point and grown for 7 days in their respective conditions. After 7 days, cells were fully trypsinized using 0.25% Trypsin-EDTA (Gibco, 25200-056) and gently scraped to collect all adherent cells. Afterwards, cells were plated in 100mm dishes at a density of 3.0 x 10^5^, 2.0 x 10^5^, and 1.5 x 10^5^ for days 3, 5, and 7 respectively under normoxic conditions. For hypoxic conditions, cells were plated in 100mm dishes at a density of 1.1 x 10^5^, 0.7 x 10^5^, and 0.35 x 10^5^ for days 3, 5, and 7 respectively. Stated densities were chosen so that cells were 80% confluent at the time of harvest.

### Flow cytometry

For flow cytometry, BMSCs were cultured as describe previously. At the indicated end points, BMSCs were harvested and stained with the LIVE/DEAD Fixable Near-IR Dead Cell Stain Kit (Invitrogen, L10119) for 30 minutes at room temperature. For antigen staining, antibodies were used at the following dilutions: CD45-PE (1:250, Invitrogen, 12-0451-81), CD45-PerCP-Cyanine5.5 (1:250, Invitrogen, 45-0451-82), PDGFRα-APC (1:250, Invitrogen, 17-1401-81). A complete list of antibodies can be found in Table S1. Cells were stained with antibodies at room temperature for 30 minutes. Flow cytometry was performed on a BD FACSCanto II (BD Biosciences, 338962) and analyzed using FlowJo software (version 10.6.1). Compensation was conducted using OneComp eBeads Compensation Beads (Invitrogen, 01-1111-41) and ArC Amine Reactive Compensation Beads (Invitrogen, A10346) according to manufacturer’s instruction using matching fluorophore-conjugated antibodies to the antibodies used for each instance. Live cells were gated for lack of anime reactive dye fluorescence. Subsequent gating was based on fluorescence-minus-one and unstained controls.

### EdU staining to assess proliferation

Proliferation was assessed through incorporation of 5-ethynyl-2’-deoxyuridine (EdU) using the Click-iT EdU Alexa Fluor 488 Flow Cytometry Assay Kit (Invitrogen, C10425). One day prior to harvest, cells were incubated with 50mM EdU for 24 hours to assess for changes in proliferation. At the time of harvest, cells were trypsinized and dishes were gently scraped to collect all adherent cells. After harvest, cells were fixed with 4% paraformaldehyde, permeabilized, and incorporated EdU was detected by a click reaction using a fluorescent Alexa Fluor 488 dye according to the manufacturer’s protocol. Following EdU detection, cells were stained with additional antibodies and subsequently analyzed with flow cytometry using the parameters described above.

### Terminal deoxynucleotidyl transferase dUTP nick end labeling (TUNEL) staining to assess apoptosis

Apoptosis was assessed through the incorporation of BrdUTP at the sites of DNA damage and detected with an Alexa Fluor 488 dye–labeled anti-BrdU antibody using the APO-BrdU TUNEL Assay Kit (Invitrogen, A23210). At the desired endpoints, cells were trypsinized and dishes were gently scraped to collect all adherent cells. Following harvest, cells were treated according to the manufacturer’s instructions for the detection of TUNEL-positive cells. Briefly, cells were fixed with 1% paraformaldehyde, permeabilized in ice cold 70% ethanol at −20°C for 60 minutes, and incubated in a mixture of TdT and BrdUTP for DNA labeling for 60 mins in a 37°C water bath. Afterward, cells were incubated for 30 minutes at room temperature in an antibody staining solution containing Alexa 488 labeled anti-BrdU. During this incubation step, antibodies against markers of interest were also added at the concentrations indicated previously. Samples were analyzed within 3 hours of completion.

### Statistical Analysis

Statistical differences were determined by one-way ANOVA followed by Tukey’s post-hoc testing or unpaired Student’s t-test, as appropriate (GraphPad, Prism 7, version 7.04). Results are expressed as mean ± SEM. p<0.05 was considered significant.

## SUPPLEMENTAL MATERIALS

**Supplemental Figure 1.**
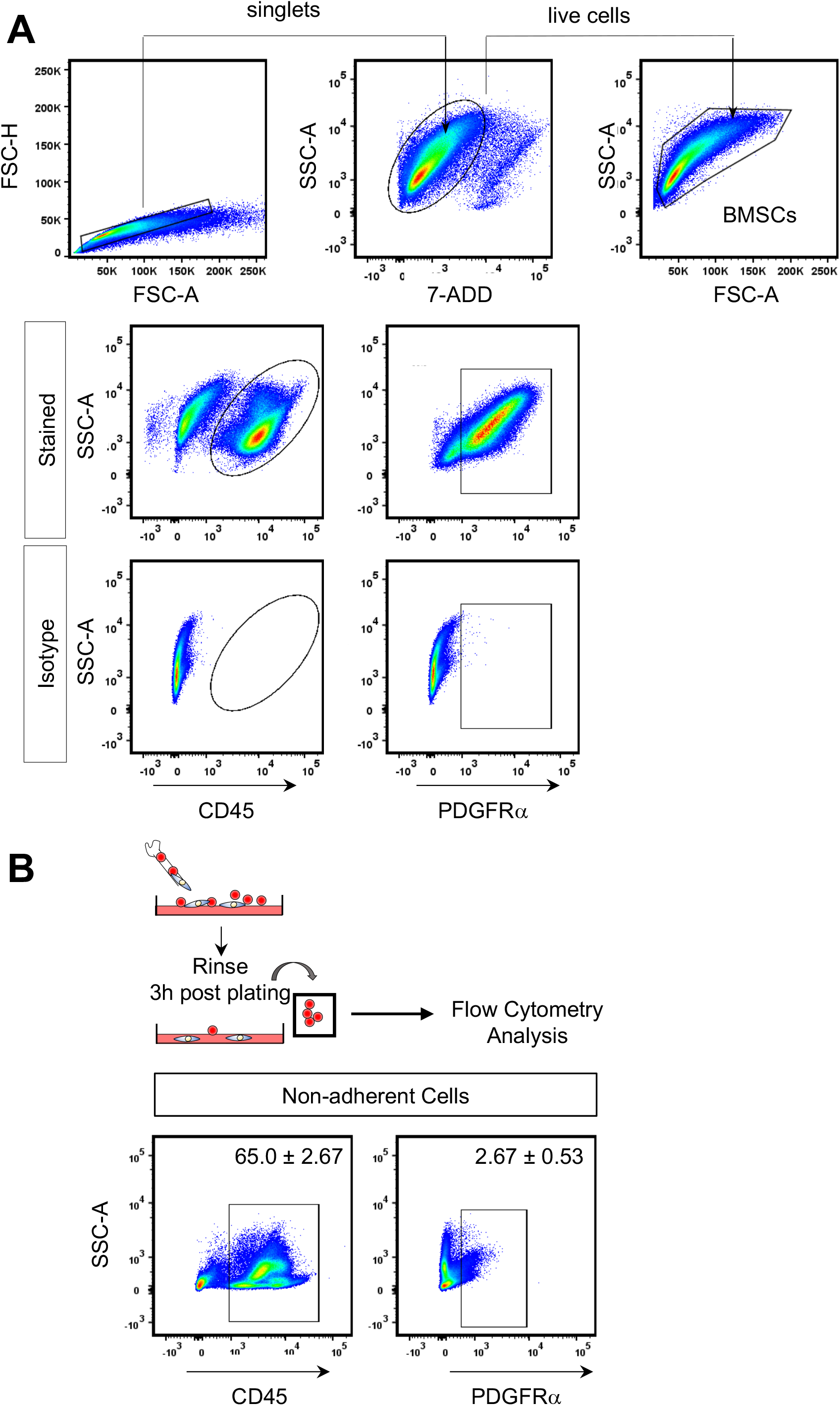
**A)** Gating strategy used for flow cytometry analysis to demark live BMSCs, CD45^+^ and PDGFRα^+^ cells staining in WT BMSC. **B)** Flow analysis for CD45^+^ and PDGFRα^+^ in non-adherent cells isolated 3 hours after plating.

**Supplemental Figure 2.**
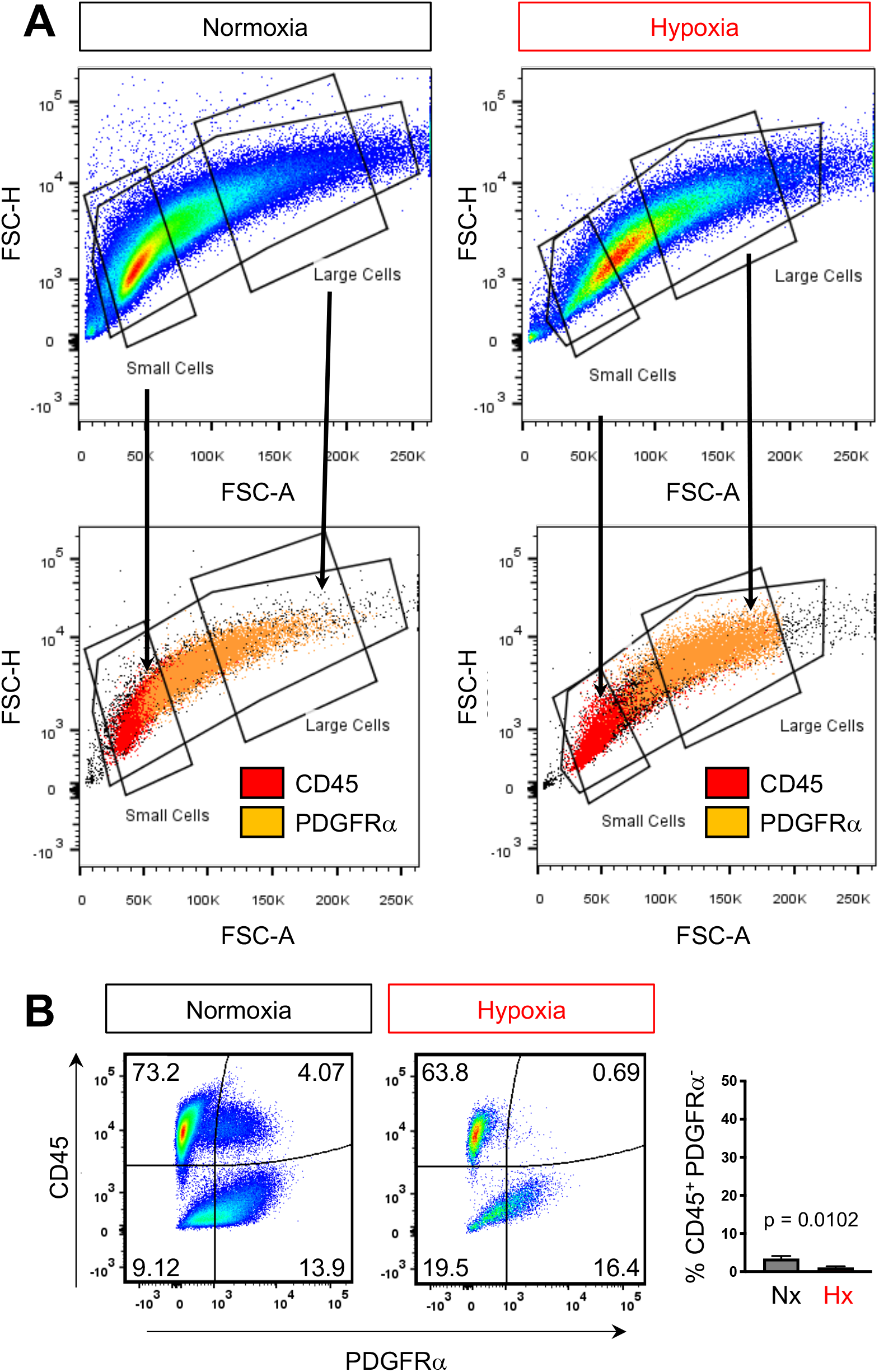
**A)** Flow analysis of BMSCs stained for CD45 and PDGFRα gated based on size.

**Supplemental Table 1.**
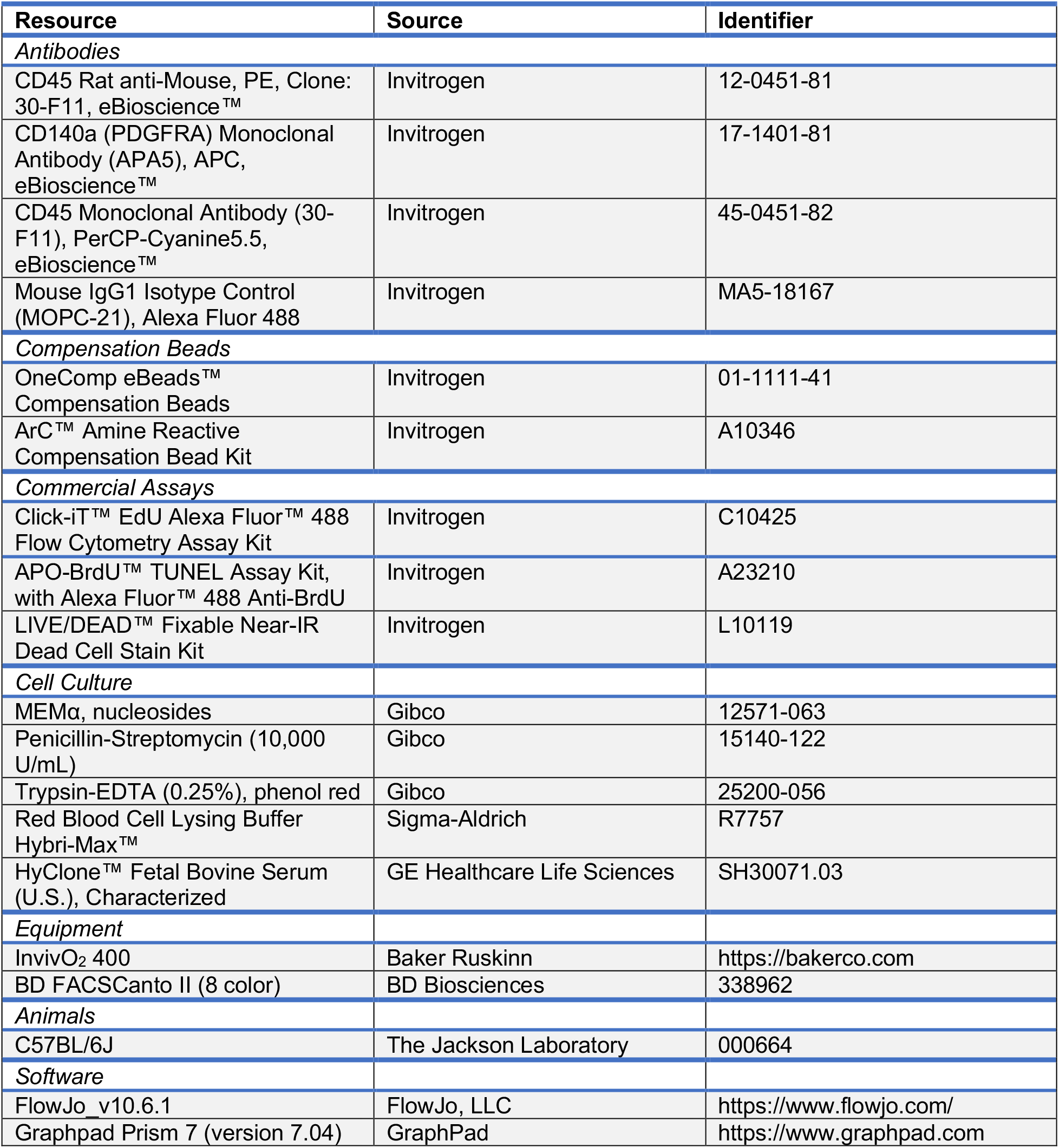
Key Resources

